# The role of human intraparietal sulcus in evidence accumulation revealed by EEG and model-informed fMRI

**DOI:** 10.1101/2025.02.05.636628

**Authors:** Jaeger Wongtrakun, Shou-Han Zhou, Redmond G. O’Connell, Trevor T.-J. Chong, Mark A. Bellgrove, James P. Coxon

## Abstract

Sequential sampling models propose that the repeated sampling of sensory information is a fundamental component of perceptual decision-making. Electroencephalographic investigations in humans have demonstrated motor-independent representations of evidence accumulation, but such observations have seldom been made in neuroimaging studies exploring the neuroanatomical origins of evidence accumulation. Here, we aimed to reveal the neuroanatomical locus of sensory evidence accumulation in the human brain by regressing an electrophysiological marker of evidence accumulation (centroparietal positivity, CPP) against changes in blood oxygen level-dependent (BOLD) signal during perceptual decision-making. Our cross-modal imaging approach revealed a cluster within left intraparietal sulcus (IPS), located within putative lateral intraparietal area (region hIP3), for which BOLD signals scaled in relation to the slope of the CPP. Furthermore, the drift rate parameter of a drift diffusion model parametrically modulated BOLD activity within an overlapping region of left IPS. In contrast, parametric modulation by reaction time revealed a distributed fronto-parietal network, demonstrating the utility of our approach for isolating a discrete neuroanatomical locus of evidence accumulation. Together, our findings provide strong support for intraparietal sulcus involvement in the accumulation of sensory evidence during human perceptual decision-making.

## 1 Introduction

Perceptual decision-making involves translating sensory information into behaviour. For example, when crossing a busy road, one must gather information about the surroundings to determine when to cross safely. Sequential sampling frameworks propose that decisions are made when evidence accumulates to a threshold to inform the choice (Ratcliff & McKoon, 2008). Remarkably, the functional neuroanatomy of this accumulation process in humans remains unclear.

The neural signatures of evidence accumulation have been a topic of investigation since accumulation-to-bound dynamics were first described in single neurons of the macaque brain (Shadlen & Newsome, 1996). Such dynamics have been observed in dorsolateral pre-frontal cortex, frontal eye fields, striatum (Kim & Shadlen, 1999; Ding & Gold, 2012ab; Heitz & Schall, 2012; Mante et al., 2013), and most extensively in the lateral intraparietal area (LIP) (e.g. Shadlen & Newsome, 1996; 2001; Roitman & Shadlen, 2002, but see Katz et al., 2016). These observations align with computational models of decision-making such as the drift diffusion model (DDM), where the intake rate of sensory evidence is encapsulated by the ‘drift rate’ parameter (Voss et al., 2004). Similarly, human electroencephalography (EEG) indexes evidence accumulation through the centroparietal positivity (CPP), a signal that builds proportionally with the strength of sensory evidence and is independent of motor preparation (O’Connell et al., 2012; Kelly & O’Connell, 2013), response modality, or visual hemifield (Loughnane et al., 2016). Although the CPP has advanced our understanding of evidence accumulation, the poor spatial resolution of EEG hinders isolation of its neuroanatomical basis. Intracranial EEG offers additional insight into abstract decision areas (e.g., Gherman et al., 2024; Xie et al., 2024), but suffers from limited brain coverage. Attempts to affirm neuroanatomical relationships between monkeys and humans can, however, be enhanced when EEG is combined with functional magnetic resonance imaging (fMRI; see Philiastides & Sajda, 2007).

fMRI studies have identified human brain regions where activity might reflect evidence accumulation during perceptual decision-making (Ho et al., 2009; Liu & Pleskac, 2011; Ploran et al., 2011; Krueger et al., 2017; Morito & Murata, 2022). The intraparietal sulcus (IPS) for instance has been implicated in tracking task-relevant and task-irrelevant evidence (Ho et al., 2009; Liu & Pleskac, 2011; Ritz & Shenhav, 2024). Studies have attempted to isolate evidence accumulation by varying sensory and/or motor requirements (e.g., Filimon et al., 2013), but have introduced temporal delays between events within trials to distinguish between cognitive processes, which may have had unintended effects (e.g. working memory-related brain activity) (Ruge et al., 2009). Consequently, there is significant variability in identified regions. The CPP is a domain-general evidence accumulation signal that can be observed across a range of different motor and sensory modalities (O’Connell et al., 2012). This unique property renders the CPP ideal for integration with fMRI to determine regions involved in sensory evidence accumulation during perceptual decision-making.

Here, we employed a multimodal approach, regressing electrophysiological (CPP slope), computational modelling parameters (DDM drift rate), and behaviour (reaction time) against BOLD signals acquired during perceptual decision-making. Across both an EEG and a subsequent fMRI session, participants performed a variation of the random dot-motion task that varied sensory feature, visual hemifield, and motor requirements (Loughnane et al., 2016; Zhou et al., 2021). We expected parametric modulation by reaction time would identify a broad fronto-parietal network (Brosnan et al., 2020) but that our specific evidence accumulation metrics would result in more spatially constrained activation. Deriving predictions from commonly reported regions in humans (Ho et al., 2009; Liu & Pleskac, 2011) and macaques (Shadlen & Newsome, 1996; 2001; Roitman & Shadlen, 2002), we hypothesised that BOLD signal changes within the human homologue of LIP within the IPS - region hIP3 (Grefkes & Fink, 2005; Richter et al., 2019) - would covary with the CPP slope across participants (greater BOLD associated with shallower CPP slope, indicating prolonged evidence accumulation). Further, consistent with resting-state fMRI connectivity (Brosnan et al., 2020), we anticipated that CPP slope would correlate with activation in secondary motor regions. Finally, we expected to see evidence of parametric modulation of BOLD activity by drift rate (i.e. within-subjects), and that there would be overlapping activity with brain regions isolated by the CPP slope regression, providing converging evidence across neurophysiology and computational modelling for the locus of sensory evidence accumulation in humans.

## 2 Methods

### 2.1 Participants

We tested 40 right-handed adults (22 females; age mean ± standard deviation [SD], 25.48 ± 6.00 years). The participants had normal or corrected-to-normal vision and no history of neurological or psychiatric disease. Participants gave their written informed consent and were tested at Monash University, Australia, in accordance with the experimental protocol approved by the Monash University Human Research Ethics Committee (#17863).

### 2.2 Experiment Setup

Participants completed the random dot motion (RDM) task (described below) while undergoing EEG, and subsequently fMRI, in separate sessions at least 24 hours apart. In the EEG session (Session 1), participants sat in a darkened room 70cm from a 27-inch LCD monitor (Dell S2716DG, running at 120Hz and 1024×768 resolution) with their head stabilised by a chin rest. The index and middle fingers of their right hand rested on the left and right buttons of a computer mouse, from which responses were registered by a button press. Eye movements were monitored using an eye tracker (DM890, Eyelink 1000 Plus Version 5.09; SR Research Ltd/SMI). Continuous EEG was acquired from each participant using the BrainVision 64 channel EEG cap and actiCHamp amplifier (Brain Products, Germany) and digitised at 500 Hz.

For the fMRI session (Session 2), participants viewed the task via a mirror attached to the head coil. The task was displayed on a 23 inch BOLDScreen in 1920 x 1080 resolution at 60Hz. Fixation was qualitatively monitored using an eye tracker (DM890, Eyelink 1000 Plus Version 5.09; SR Research Ltd/SMI). Participants utilised their right index and middle finger to respond to task stimuli via a MR-compatible button box.

The RDM task involved monitoring two kinematogram patches of dots. In the EEG and fMRI sessions, respectively, the two patches were centred at 10° / 7.5° on either side of fixation and 4° / 3° below the horizontal meridian. Each patch subtended 8° / 6° of visual angle and comprised 150 white dots (EEG 6 x 6 pixels, fMRI 13 × 13 pixels) against a black background.

### 2.3 Experimental Task and Protocol

We used the bilateral kinematogram variant of the RDM discrimination task (Figure 1) (Loughnane et al., 2016; Zhou et al., 2021). The task structure was identical across the EEG and fMRI sessions (except for the inclusion of rest blocks during fMRI) and each task run corresponded to a factorial design. The factor Attended Modality (Attend to Motion, Attend to Colour) was manipulated at the block level (Figure 1A), whereas the factors Coherent Motion Direction (Up, Down), Colour Change (Red, Blue), Target Hemifield (Left, Right), and Distractor Congruence (Congruent Hemifield, Incongruent Hemifield) were fully counterbalanced across trials. Each run consisted of eight experimental blocks (four per Attended Modality), with each block comprising eight trials, totalling 64 trials per run. The task used in this experiment included the Distractor Congruence factor for the purpose of examining the effect of spatial congruence of target (motion) and distractor (colour) on behaviour and BOLD signal change (i.e., whether changes in coherent motion occurred in the same vs different visual hemifield to changes in colour). In this manuscript we collapse across the levels of this factor. The investigation of distractor-mediated effects will be the focus of a separate manuscript.

**Figure 1.**
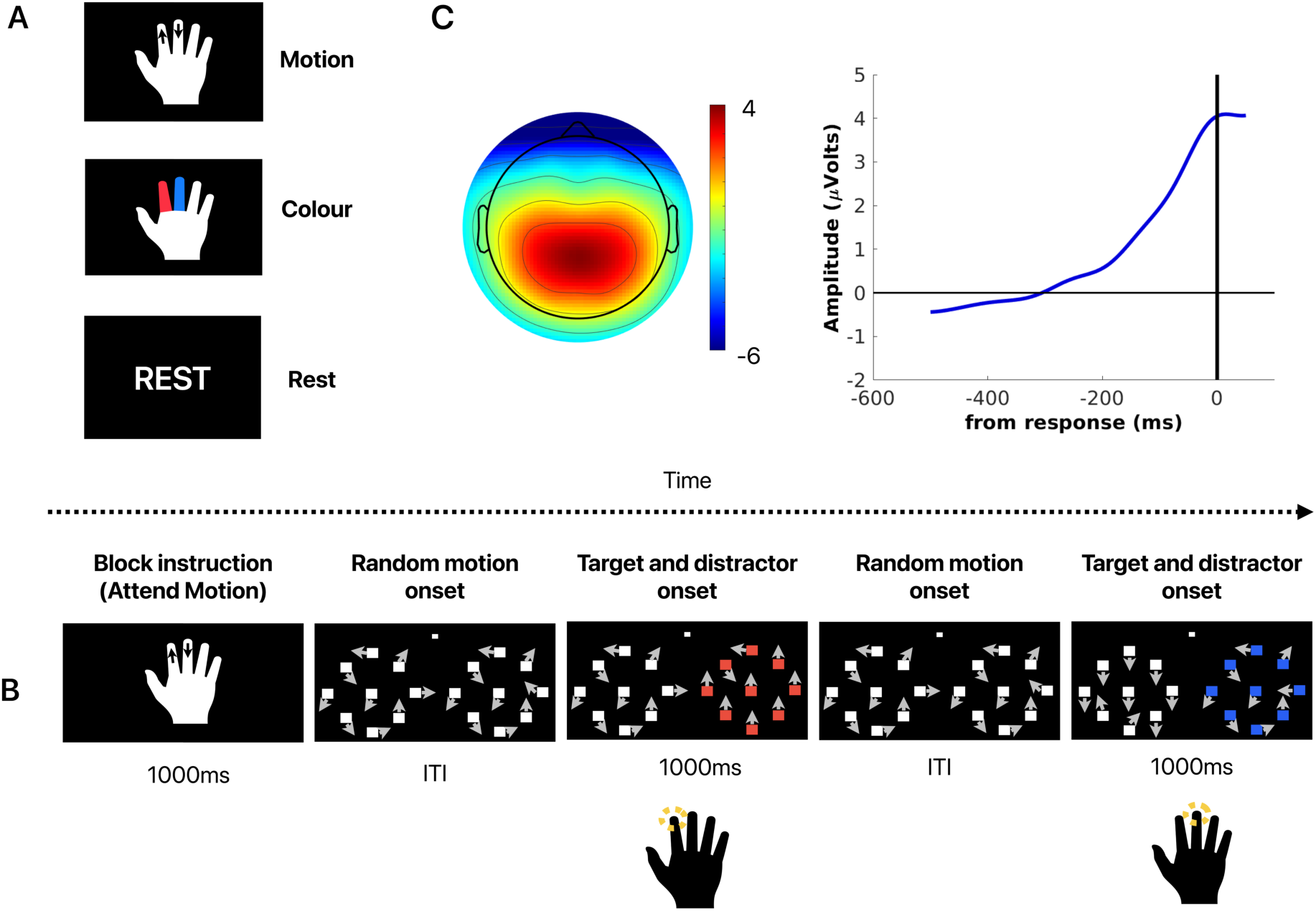
Experimental task. **A)** Instructional cues to attend to motion (arrows) or colour (colours), or neither (rest), were presented to participants at the beginning of each block of eight trials within each run. **B)** Shown are two trials of a Motion respond block with descriptions and durations of events respectively listed above and below. A cue was presented at the beginning of the block, instructing participants to attend to changes in motion. The intertrial interval (ITI) lasted between 1000 ms to 3500 ms in total, split into a fixation phase, where all dots froze for a duration between 1000ms to 2500 ms, and the random dot-motion phase, where all dots underwent random motion for between 900 ms to 1000 ms. During the first trial, at target onset a proportion of the dots in the right patch transitioned from random to upwards coherent motion for 1000ms. A change in colour also occurred, in this case all dots in the right patch also underwent a change to the colour red. The participant indicated the change in the attended modality (upwards motion) via a right index finger button press. During the second trial, at target onset a proportion of the dots in the left patch transitioned to downward motion, while the dots in the right patch underwent a change to the colour blue. The participant indicated the change in the attended modality (downward motion) via a right middle finger button press. For colour blocks, participants attended to the colour change and responded according to the cued response mapping. **C)** Topography and time course of the response-locked CPP.

Throughout the task, participants focused on a central point while monitoring two peripherally located patches of random dot stimuli (Figure 1B). The patches underwent two changes on each trial: a transition to coherent motion and a change in colour. These changes occurred in either the same patch of dots (e.g. motion and colour change both occurred in the right patch) or in opposing patches of dots (e.g. motion change occurred in the left patch, colour change occurred in the right patch). For each session (EEG, fMRI), participants completed four scanner runs. During the first two runs, participants were instructed to mentally tally the number of times they observed a change within the attended stimuli (Count runs). Participants were cued to attend to only a single change per block (a single up or down arrow; or a single blue or red colour) and at the end of counting blocks they were given a short period of time to report the number of instances of change that they had observed (e.g., five button presses to indicate having observed five changes in downwards motion). During the subsequent two Respond runs, participants immediately indicated when they observed a change in the attended stimulus via button press. Each of the two options per modality (upwards/downwards motion or blue/red colour) were assigned to a participant’s index or middle finger and held constant across both EEG and fMRI sessions. The stimulus/response mapping was fully counterbalanced across participants.

Each participant underwent a titration procedure prior to commencing their first session. The proportion of dots that transitioned from random to coherent motion was determined via a thresholding algorithm that titrated participants to achieve approximately 70% success (Taylor & Creelman, 1967).

For both EEG and fMRI sessions, participants were presented with an instructional cue for 1000 ms at the beginning of each block, indicating which modality to attend to. Each trial was separated by an intertrial interval (ITI) that was based on an exponential decay distribution with a minimum of 2000 ms, mean of 2300 ms, and a maximum of 3500 ms. The ITI consisted of an initial period of non-moving dots lasting between 1000 ms and 2500 ms, followed by pre-target random motion, in which dots were randomly positioned within each patch throughout each frame. After a random period of 900-1000 ms, a proportion of the dots in one of the patches transitioned from random to coherent upwards or downwards motion.Coincident with the onset of coherent motion (sampled from a normal distribution centred on coherent motion onset, SD = 50ms), the colour of all dots in either the same or alternate patch changed colour to blue or red. The onset times for motion and colour change were jittered in an attempt to prevent participants from using the change in colour to prime participants’ attention and thereby bias their perception (Failing & Theeuwes, 2020; see Supplementary Data). Each trial terminated 1000 ms after the onset of the attended stimulus.

In the fMRI session, task blocks (32 second duration) were interleaved with Rest blocks (24 second duration). Rest blocks involved the same visual stimuli, except that participants simply observed the display without counting or responding.

### 2.4 EEG Preprocessing

EEG data were processed using a combination of custom scripts and EEGLAB routines implemented in MATLAB based on a modified Harvard Automated Processing Pipeline for Electroencephalography (Gabard-Durnam et al., 2018; Stefanac et al., 2021). The signals were band pass filtered between 0.01 Hz and 35 Hz using a fourth-order Butterworth filter before re-referencing to the average of all channels. Channels with a high amount of noise were interpolated using a spherical spline. An autonomous independent component analysis was applied to remove EEG artifacts such as muscle contractions, meanwhile trials were excluded if fixation was broken (either by blinking or eye movement) >3° left or right of central fixation point. EEG epochs between −700 to +1880 ms around target stimulus onset were then baseline corrected to the mean amplitude over the 100ms prior to stimulus onset (i.e., from −100 to 0 ms). For the attend motion respond condition, the slope of the CPP waveform was measured from the peak electrode CPz (Steinemann et al., 2018; Newman et al., 2017; Kelly & O’Connell, 2013) and determined as the gradient of a straight line, fit from −400 to −50 ms of the response-locked CPP waveform (Figure 1C; Loughnane et al., 2016).

### 2.5 Fitting the Drift Diffusion Model

We fit a drift diffusion model to the reaction time data from the fMRI session to estimate latent parameters that summarise participants’ decision-making during the attend-motion condition. Using the HDDM package v0.6.0 (Wiecki et al., 2013), single trial response time (RT) and accuracy data were modelled as a Wiener first passage time (wfpt) process. Motion trials were excluded if RTs were less than 300 ms or greater than 1500 ms after the change in coherent motion (< 1 % of trials). These thresholds were implemented to remove pre-emptive and/or slow responses (Zhou et al., 2021). For the purpose of employing drift rate as a parametric modulator (discussed later), RTs were split into tertiles for each participant, resulting in three RT bins: fast, medium, and slow. The model specified three parameters: the drift rate, *ν*, the boundary separation, *α*, and the non-decision time, *τ*. The three RT bins (fast, medium, slow) were included as a condition in the DDM. Note that because the experimental design was factorial (conditions were equiprobable), participants were not able to anticipate the direction of the dot motion. We included a bias term in the model, assuming an equal probability for participants to perform correctly or incorrectly at the onset of the stimulus change (i.e., bias = 0.5; 50% correct, 50% incorrect). In other words, participants did not have information prior to trial onset, e.g. an attentional cue, that would bias their behaviour. We have omitted the bias term from the model equation below to reflect that bias is independent of RT.

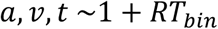

The DDM parameters for each participant were estimated using single trial reaction times by generating 100,000 samples from the joint posterior distribution of the DDM parameters using Markov Chain Monte Carlo methods (Gamerman & Lopes, 2006). To ensure convergence and minimise the effect of initial values on the posterior inference, the initial 80,000 samples were discarded as burn-in. Subsequently, the parameter estimates for each RT bin condition were used in the event-related fMRI as a drift rate parametric modulator with three levels, as described below. With the focus of this manuscript being evidence accumulation, drift rate was the parameter of interest.

### 2.6 fMRI Acquisition and Preprocessing

Imaging was performed using a Siemens Skyra 3T MRI scanner (Siemens, Erlangan, Germany), located at Monash Biomedical Imaging, Monash University, equipped with a 20-channel head coil. A T1-weighted structural image (TR = 2.3 s; TE = 2.1 ms; flip angle = 9; FOV = 240 x 256; voxel size = 1 mm, isotropic) and a gradient field map (TR = 0.447 s; TE1 = 7.38 ms; TE2 = 4.92 ms; flip angle = 60; FOV = 64 x 64; voxel size = 3.59 x 3.59 x 3 mm) were obtained for each participant. Functional MRI data was subsequently acquired using an echoplanar imaging sequence for T2* weighted BOLD contrast (TR = 2.43 s; TE = 30 ms; flip angle = 90; FOV = 532 x 532; voxel size = 3.026 x 3.026 x 3 mm, descending slice acquisition).

Preprocessing of the fMRI data was performed using fMRIPrep version 21.0.1 (Esteban et al., 2019). First, a reference volume and its skull-stripped version were generated. BOLD images were realigned to the reference EPI, unwarped using the acquired fieldmap image, and slice-time corrected to the middle slice. The EPI volumes were then co-registered with six degrees of freedom to the T1w reference. Next, the T1w reference was spatially normalised through nonlinear registration to MNI152NLin2009cAsym space and the EPI volumes were resampled into this MNI standard space. Denoising was then performed via automatic removal of motion artifacts using independent component analysis (ICA-AROMA, Pruim et al. 2015) after removal of non-steady state volumes and spatial smoothing with an isotropic, Gaussian kernel of 6mm FWHM (full-width half-maximum).

### 2.7 fMRI Model Specification and Analysis

Preprocessed functional imaging data were used as input for the first-level general linear models (GLM) of the four runs (two Count, two Respond) via SPM12 (https://fil.ion.ucl.ac.uk/spm/software/spm12/). For Count and Respond runs, eight separate conditions were specified (Count Motion Left, Count Motion Right, Respond Motion Left, Respond Motion Right, Count Colour Left, Count Colour Right, Respond Colour Left, Respond Colour Right). To obtain sufficient trials with correct responses per condition for model estimation we collapsed the factors Coherent Motion Direction (Up, Down) and Colour Change (Red, Blue).

Event-related designs were specified for each participant with the conditions outlined above, with Rest as an implicit baseline. Events pertaining to instructional cues and error trials were added as regressors of no interest, meaning they were specified in the model but given no weighting when creating contrasts. The onset of coherent motion (attend to motion blocks) or colour (attend to colour blocks) was modelled as the event of interest with a stick function (i.e. duration of 0 seconds). Separate event-related models were estimated as described above, with the addition of i) RT and ii) drift rate iii) the decision threshold and iv) non-decision time from the DDM as the parametric modulator in the Respond runs.

Several confounding time-series were calculated based on the preprocessed BOLD signals and also included as regressors of no interest: six head-motion parameters, framewise displacement, DVARS, and three region-wise global signals. The three global signals were extracted from cerebrospinal fluid, white matter, and a whole-brain mask. Additionally, a set of physiological regressors were extracted to allow for component-based noise correction. Principal components were estimated after high-pass filtering the preprocessed BOLD time-series (using a discrete cosine filter with 128s cut-off) for the two CompCor variants: temporal (tCompCor) and anatomical (aCompCor).

Motion change was the primary modality of interest, with contrasts specified separately for Count and Respond runs as follows (six total): Attend Motion > Rest; Attend Motion Left > Rest, Attend Motion Right > Rest. For the Respond contrasts only, an additional set of contrasts were specified for the parametric modulator (three for each, twelve total). These contrast images resulting from the first-level analyses were used as input for second-level group analyses. In addition to specifying one-sample t-tests (voxel level family-wise error corrected, FWE < .05), a conjunction analysis (Nichols et al., 2005) was specified for Motion to determine which regions remained active when motor execution and attentional selection processes were controlled. Contrasts for Respond and Count runs from each visual hemifield were included (i.e., Attend Motion Count Left ∩ Attend Motion Count Right ∩ Attend Motion Respond Left ∩ Attend Motion Respond Right), each thresholded at the cluster level (height threshold t > 2.71, *p* < .005, FWE < .05). The resulting conjunction image was then applied as an implicit mask for the Attend Motion Respond > Rest contrast to determine peak MNI coordinates within the conjunction.

Next, three regression models were specified: i) The *response*-locked CPP slope signal obtained from each participant during the EEG session respond runs was regressed against the Attend Motion Respond > Rest fMRI contrast, masked by the conjunction image, ii) The *stimulus*-locked CPP slope signal from count runs was regressed against the Attend Motion Count > Rest fMRI contrast, iii) The mean RT values from the fMRI session per participant were regressed against the Attend Motion Respond > Rest fMRI contrast. Parametric modulation and regression analyses were corrected for multiple comparisons at the cluster level (height threshold t > 2.71, *p* < .005, FWE < .05). Using the Jülich histological atlas (Amunts et al., 2020), a bilateral anatomical ROI of area hIP3 was created. hIP3 is the putative human homologue of the macaque lateral intraparietal area (Grefkes et al., 2005; Richter et al., 2019). Small volume correction using this ROI was then applied to the parametric modulator contrast.

### 2.8 Statistical Analysis of Behavioural Data

Analyses of behavioural data were conducted using JASP Version 0.17.2.1 (JASP Team, 2023). Mean RTs and accuracy were calculated for the Motion respond condition for the two sessions per participant. To determine feasibility of utilising data from the EEG session in second-level fMRI regression models, RT and accuracy across the EEG and fMRI sessions were compared with a paired samples t-test and correlation analysis.

## 3 Results

### 3.1 Behavioural Results

There was a strong positive relationship between the response times acquired during the EEG and fMRI sessions (r = 0.62, *p* < .001, Figure 2), indicating that the CPP slope obtained during EEG could be appropriately applied in a subsequent fMRI regression analysis. Response times were significantly longer in the fMRI session (Mean RT = 743ms, SD = 82ms) compared to the EEG session (Mean RT = 665ms, SD = 69ms, t(39) = 7.35, *p* < .001. Similarly, accuracy was lower in the fMRI session (Mean accuracy = 86%, SD = 8%) than the EEG session (Mean accuracy = 92%, SD = 6%, t(39) = 4.369, *p* < .001). These session effects are likely due to the additional challenges of presenting stimuli and recording responses in the MRI scanner.

**Figure 2.**
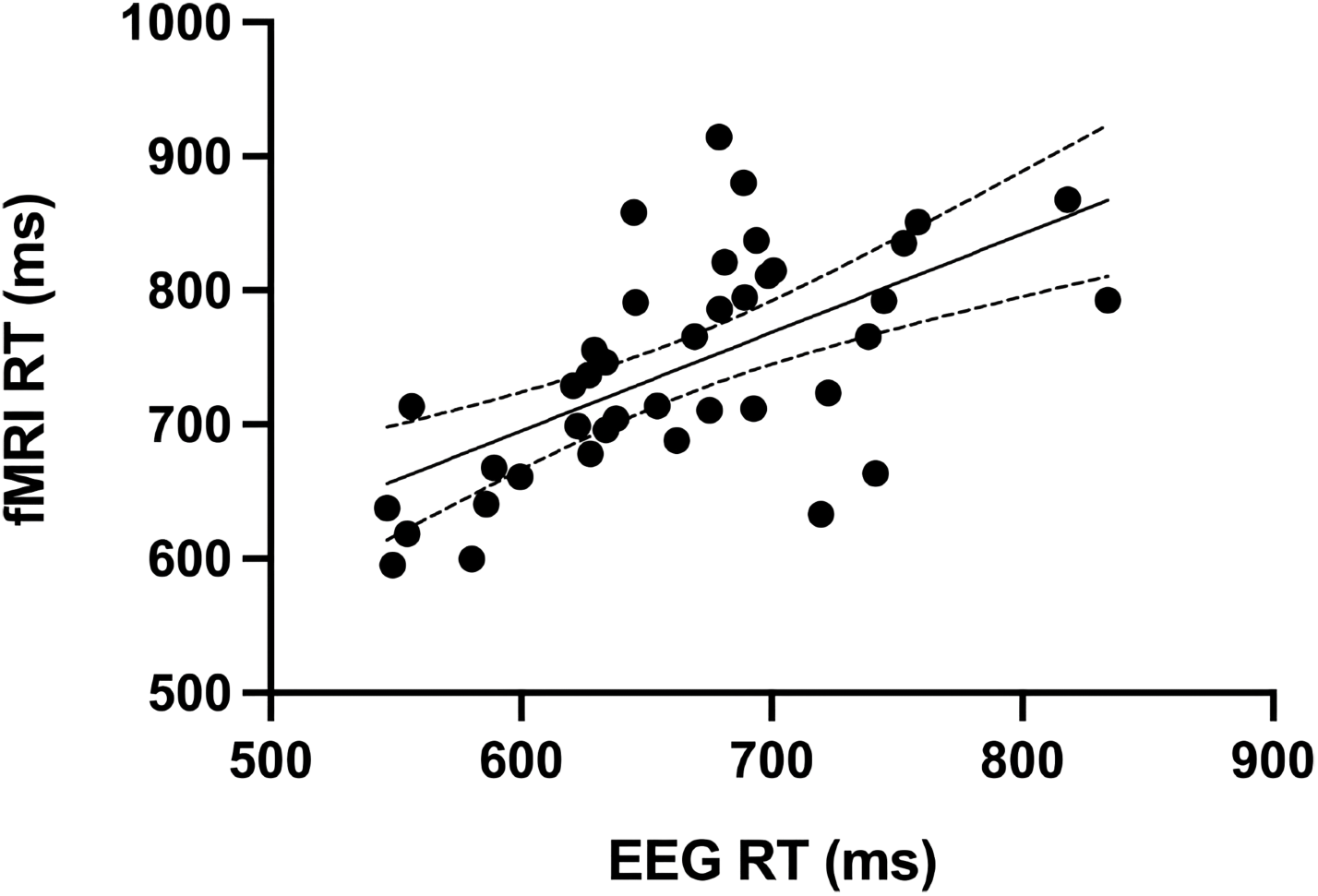
Response times for the EEG and fMRI sessions were positively correlated (r = 0.62, *p* < .001).

### 3.2 Functional Magnetic Resonance Imaging

The experiment was primarily designed to reveal the functional neuroanatomical locus of sensory evidence accumulation. First, a conjunction analysis was conducted for the purpose of isolating evidence accumulation from attentional selection and motor processes, consisting of the following conditions: Attend Motion Count Left ∩ Attend Motion Count Right ∩ Attend Motion Respond Left ∩ Attend Motion Respond Right. The conjunction analysis revealed activations bilaterally in pre supplementary motor area (preSMA), premotor areas, anterior insula, IPS, caudate, and putamen (Figure 3A, see Supplementary Data, Table 1 for coordinates and Supplementary Data, Figure 1 for the input images). This identified candidate regions that are potentially involved in sensory evidence accumulation.

**Figure 3.**
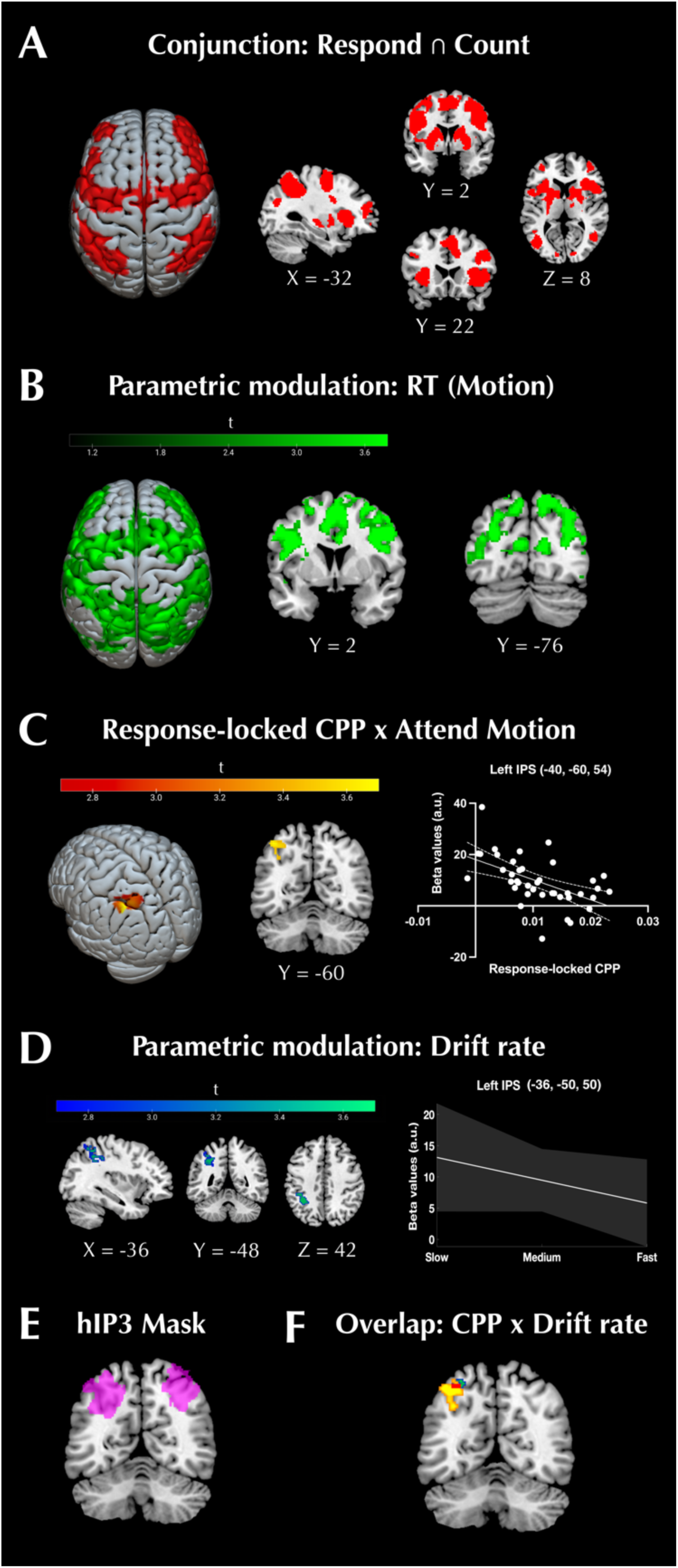
Evidence supporting the left IPS as a neuroanatomical locus of the CPP signal. A) Conjunction analysis. Activations common to Respond and Count conditions across left and right visual hemifields, which are thought to have been isolated from execution- and attentional-related processes (see Methods). B) RT as a parametric modulator revealed a diffuse activation network. C) Response-locked CPP slope as a regressor in a second level model using the Attend Motion Respond > Rest contrast images for each participant. Shallower CPP slopes were associated with greater BOLD signal in the left IPS after small volume correction. D) Drift rate as a parametric modulator revealed an inverse relationship between BOLD activation in left IPS with drift rate (i.e., smaller drift rate values correlated with greater BOLD signal). E) Area hIP3, i.e., putative LIP, used as a bilateral mask for small volume correction for these analyses. F) A binarised, uncorrected (*p* < .005) overlay of response-locked CPP (yellow), drift rate (blue-green), and the overlapping voxels (red), supporting a region of the left IPS as being central in the process of accumulating sensory evidence.

Next, we examined how RT covaried with BOLD signal within-subjects when participants attended to motion changes during perceptual decision-making. At the whole-brain level (Figure 3B), this analysis revealed a diffuse bilateral fronto-parietal activation network when participants attended to Motion, with slower RTs being correlated with greater BOLD activations. We also explored how RT covaried with BOLD signal within-subjects when participants attended to changes in colour (Supplementary Data, Figure 2). This analysis revealed a set of clusters that centred around regions of the ventral attention and salience networks (Seeley et al., 2007; Vossel et al., 2014; Chand & Dhamala, 2016). The similarity between the conjunction (Figure 3A) and RT parametric modulation (Figure 3B) indicates that the regions commonly active across conditions also scaled their activity with the speed of perceptual decisions. However, it is unlikely that all regions are important for the sensory evidence accumulation process.

To interrogate the neural correlates of sensory evidence accumulation more specifically, a second level model was specified using the Attend Motion Respond > Rest contrast images (masked by the conjunction) and regressed against response-locked CPP slope values, our electrophysiological measure of evidence accumulation. This revealed a single cluster in the left IPS (MNI −40, −60, 54) (Figure 3C; see Supplementary Data, Figure 3 for whole-brain analysis). In contrast, we found that the same contrast regressed against RT produced a single cluster in right middle frontal gyrus (Supplementary Data, Figure 3), which has been reported by some as a convergence point for dorsal and ventral attention networks (Japee et al., 2015). An additional between-subjects regression analysis was specified using the Attend Motion Count > Rest contrast images and the stimulus-locked CPP slope. A cluster was identified within left anterior IPS with a liberal height threshold (Supplementary Data, Figure 5; MNI −32, −38, 38, *p* < .05, t > 1.69, uncorrected). Collectively, the analyses reported thus far highlight the utility of using the (response-locked) CPP slope to identify the locus of sensory evidence accumulation in the left IPS.

To determine the CPP’s relationship with previously reported coordinates implicated in evidence accumulation in inferior frontal cortex, insula, and supplementary motor cortex (Ho et al., 2009; Kayser et al., 2010; Filimon et al., 2013), we created spherical ROIs (10mm radius) and extracted beta values to examine each region’s relationship with the CPP slope (Supplementary Data, Figure 6). All correlations were non-significant (*p* > .05), with the exception of the left inferior frontal sulcus from Filimon et al. (2013) (r = −.4, *p* = 0.01). This suggests that human sensory evidence accumulation may not be as spatially distributed as previously thought, and may primarily take place in left IPS (Figure 3).

Next, we set out to provide complementary evidence with model-based fMRI, by incorporating a computational parameter that indicates the intake rate of sensory information i.e., drift rate (Voss et al., 2004). The drift rate parameter was included as a within-subjects parametric modulator in the first-level GLM. At the whole brain level (Supplementary Data, Figure 7), we identified a similar cluster within left IPS, where BOLD activity increased as a function of drift rate, i.e., the lower the drift rate, the greater the BOLD activation. We applied small volume correction with the bilateral anatomical ROI of area hIP3 (Figure 3E), again revealing a single significant cluster within left IPS (Figure 3D; MNI −36, −50, 50, *p* < .005, t > 2.69, FWE < .05 cluster-wise). In addition, we ran analyses using decision threshold and non-decision time parameters as the parametric modulator instead of drift rate. No voxels survived the cluster level inference for decision threshold (FWE > .05). However, parametric modulation by non-decision time (Supplementary Data, Figure 8) revealed a similar activation to the parametric modulation using RT (Figure 3B), suggesting these parameters were highly correlated, which we confirmed via correlational analyses (fast: r = .94; medium: r = .98; slow: r = .94). Overall, these analyses indicate that both electrophysiological (CPP slope) and computational (drift rate) metrics of sensory evidence accumulation are associated with the BOLD signal in left IPS, whereas RT and non-decision time exhibit a spatially unspecific association.

Finally, we evaluated the similarity between the activation for the response-locked CPP slope regression and the parametric modulation of drift rate by overlaying the contrast images, and we observed overlap within the left IPS (Figure 3F). Additionally, we confirmed that this cluster fell within the common network activated across contrasts (conjunction analysis, see Methods and Figure 3A). Collectively, these results provide evidence suggesting the left IPS as the neuroanatomical origin of the CPP signal.

## 4 Discussion

Here, we aimed to elucidate the neuroanatomical locus of sensory evidence accumulation in the human brain by regressing CPP slope and DDM drift rate parameter values against changes in BOLD signal during perceptual decision-making. Our multimodal imaging approach, in which we controlled for the confounding processes of feature-selective sensory areas (attentional selection) and motor preparation, revealed a cluster within left IPS for which BOLD signal scaled in relation to the slope of the CPP. In addition, we showed that a computational parameter indexing rate of sensory evidence intake (drift rate) parametrically modulated BOLD activity (within-subjects) within an overlapping region of the left IPS. In contrast to the spatially specific IPS activation associated with sensory evidence accumulation metrics, reaction time and the DDM non-decision time parameter showed spatially diffuse association with BOLD activity. Our findings thus strongly suggest involvement of the left IPS in the accumulation of sensory evidence during perceptual decision-making.

### 4.1 CPP Slope Regression

Here, we present the first fMRI-based evidence for the origin of the CPP signal during perceptual decision-making. EEG and magnetoencephalography (MEG) have been used to explore possible regions contributing to the CPP signal. Gherman et al. (2024) used intracranial EEG from epilepsy patients to characterise evidence-dependent, but motor response independent, build-up signals while participants performed a perceptual decision-making task. In the pooled data, abstract accumulation signals were observed bilaterally within prefrontal, parietal, inferior temporal, and anterior insular areas. Further, the subset of participants with contact points in parietal regions exhibited CPP-like signalling, scaling with the strength of sensory evidence and peaking at the time of the response. Although it is important to acknowledge that the relationship between scalp/intracranial EEG and fMRI is complex (see Ebrahiminia et al., 2022), these findings are consistent with our findings of a largely motor independent network involved in evidence accumulation (population-level inference, random effects analysis, see Figure 3A). Others have used MEG source reconstruction to show that the IPS tracks the evolving action plan as well as the accumulation of task-relevant evidence (Wilming et al., 2020). Overall, the above findings demonstrate a plausible link between parietal areas and evidence accumulation signals, a pattern we confirmed in our imaging data.

To identify the neuroanatomical locus of the CPP, we regressed its slope against BOLD activity to reveal a significant cluster within left IPS. Parietal activity is consistent with previous work suggesting left parietal areas to be the source of the CPP (Herding et al., 2019). Acknowledging the limitations of EEG source localisation (see Michel & He, 2019), Herding and colleagues localised the CPP from a vibrotactile-based perceptual decision-making task to the left superior parietal lobule. Our findings indicate that shallower CPP slopes during visual evidence accumulation were linked to greater BOLD activations within the medial wall of left IPS. Discrepancy between the present findings and those of Herding and colleagues could relate to sensory modality, or uncertainty in EEG source localisation. Regardless, the findings converge to identify a probable left parietal contribution to the formation of the CPP signal.

An important consideration regarding investigations of evidence accumulation is the impact of motor activity. Studies that have implicated the IPS (Ploran et al., 2007, 2011; Ho et al., 2009; Liu & Pleskac, 2011) and LIP (Shadlen & Newsome, 1996; 2001; Roitman & Shadlen, 2002) in evidence accumulation often utilise motor task responses. Recent work suggests that motor-independent choice signalling can occur even with action-linked decisions (Wu et al., 2019; Sandhaeger et al., 2023). We included a counting condition in an attempt to partition the impact of motor-related processing. Using a conjunction analysis of the Respond and Count conditions as an explicit mask (Figure 3A), we showed that left IPS remained active when motor activity was uncoupled from the decision (Figure 3C). Left IPS specifically has been demonstrated to carry perceptual choice-related information independent of motor-related processing during fMRI (Wu et al., 2021).

To substantiate the left IPS as the neuroanatomical locus of the CPP, we also looked at whether the activations were related to RT. Our parametric modulation of RT revealed a highly diffuse network. When we included average RT in a between-subjects regression model (Supplementary Data, Figure 4), we found a single cluster in the right middle frontal gyrus. This region is thought to be a nexus point for the ventral and dorsal attention networks, and plays a critical role in the reorienting of attention (Fox et al., 2006; Japee et al., 2015). This cluster was not present within our between-subjects response-locked CPP regression, suggesting that the CPP indexes a distinct process from that captured by RT during perceptual decision-making. These findings further support functional differences between CPP and RT.

In the present study, participants were required to apply top-down attentional control to discriminate between colour and motion changes and select the appropriate visual hemifield. Previous work has identified IPS regions within area hIP3 that show a parametric response to both attended to and ignored stimuli (Kayser et al., 2010), proposing that some parts of the IPS regulate attentional control, whereas others facilitate processes such as evidence accumulation. This notion aligns with recent work showing that the IPS holds diverse roles in attentional processing (Ritz & Shenhav, 2024), similar to neurons within LIP (e.g., Meister et al., 2013; Seideman et al., 2022). Specifically, the IPS encodes both target and distractor information (Ritz & Shenhav, 2024). CPP dynamics have been reported to be influenced by target selection signals (Loughnane et al., 2016), and we have previously shown that the presence of distractors can impact CPP slope amplitude (Zhou et al., 2021). There appears to be similarity between functions of the IPS and factors impacting CPP dynamics. Given the role of attention as a filter for task-irrelevant information during perceptual decision-making (Rangelov & Mattingley, 2020), this speaks to a core function of the left IPS when gathering sensory evidence, although future work examining distractor processing within left IPS is required.

### 4.2 Using computational models to support the anatomical origins of evidence accumulation

We took a novel approach in examining evidence accumulation by parametrically modulating BOLD activity within-subjects using the DDM drift rate parameter. The CPP indexes evidence accumulation whereas the drift rate represents the average intake rate of sensory evidence (O’Connell et al., 2012; Voss et al., 2004). Other fMRI studies have systematically manipulated the degree of coherence of motion stimuli, and by proxy the rate of evidence accumulation, confirming the effect of coherence by fitting a DDM to participant responses (Ho et al., 2009; Liu & Pleskac, 2011). Our examination of evidence accumulation differed to these studies in that we used a DDM to directly determine whether BOLD activity was modulated as a function of the drift rate parameter.

We observed that when drift rate values were lower (indicative of longer evidence accumulation), BOLD signal within left IPS increased. We also observed additional activations in dorsal anterior cingulate cortex and preSMA (Supplementary Data, Figure 7). Although these regions were not found within our whole-brain analysis for CPP slope (Supplementary Data, Figure 3), these activations are consistent with studies of cognitive control (Ritz & Shenhav, 2024) and determining decision thresholds (Forstmann et al., 2008) – which were likely to occur during our task. It is unclear why these regions were absent from the CPP whole-brain analysis, although it is possible that, whereas the CPP is independent of response modality (O’Connell et al., 2012; Twomey et al., 2016), drift rate can encompass response preparation processes (Dmochowski & Norcia, 2015). We acknowledge that drift rate and CPP are not identical indices of evidence accumulation; however, parametric modulation of BOLD signal via drift rate produced activation that converged with our other findings for the left IPS being important for evidence accumulation.

### 4.3 Limitations

EEG and fMRI data were collected in separate sessions. While it is technically possible to acquire EEG concurrently with fMRI, this is not without its challenges and can impact data quality. Notwithstanding, session effects could have added experimental noise, but if anything, this should have *weakened* our ability to identify the observed relationship between CPP slope and BOLD activation in left IPS. This finding was supported by convergent evidence from computational drift diffusion modelling, though establishing causal relationships between left IPS and evidence accumulation using techniques such as transcranial magnetic stimulation will be important. Additionally, it will be necessary for future work to explore whether other factors connected to CPP signalling, such as confidence (Boldt et al., 2019; Grogan et al., 2023; Ko et al., 2024; Dou et al., 2024) and urgency (Yau et al., 2020; 2021), also share a relationship with the left IPS during perceptual decision-making. Lastly, given the CPP’s relationship with distractors (Zhou et al., 2021), future research will be required to determine the effect of distractor processing within left IPS when making perceptual decisions.

### 4.4 Conclusion

In summary, this fMRI study elucidates a probable neuroanatomical locus of the CPP signal in the left IPS and supports previous work in identifying regions involved in facilitating evidence accumulation. In line with work in non-human primates and studies of cross species homology, our findings support the hIP3 region as the putative LIP region in humans and its role in sensory evidence accumulation. Our findings clarify the extent to which parietal regions, and specifically the IPS, contribute to the process of evidence accumulation during perceptual decision-making.

## Data and code availability

Data and code available upon reasonable request to Corresponding Author.

## Author contributions

J.W.: Data acquisition, analysis, manuscript writing, editing.

S-H.Z.: Data acquisition, analysis, manuscript writing, review, and editing.

R.G.O.: Manuscript review and editing, funding acquisition.

T.T-J.C.: Manuscript review and editing, funding acquisition.

M.A.B.: Manuscript review and editing, funding acquisition.

J.P.C.: Manuscript review and editing, funding acquisition.

## Acknowledgements

The authors acknowledge the facilities and scientific and technical assistance of the National Imaging Facility (NIF), a National Collaborative Research Infrastructure Strategy (NCRIS) capability at Monash Biomedical Imaging (MBI), a Technology Research Platform at Monash University.

Sarah Wallis (data collection).

