## Supplementary Data for "The role of human intraparietal sulcus in evidence accumulation revealed by EEG and model-informed fMRI"

**Table 1.** MNI coordinates of activation peaks for the Attend Motion Respond > Rest contrast spatially constrained by the conjunction analysis.

| Region | Left hemisphere | | | Right hemisphere | | |
|---|---|---|---|---|---|---|
| | **MNI (x,y,z)** | **t-value** | **p-value** | **MNI (x,y,z)** | **t-value** | **p-value** |
| **PreSMA** | -4, 0, 54 | 10.85 | < .001 | 2, 6, 58 | 8.43 | < .001 |
| **Premotor areas** | -54, 10, 34 | 11.00 | < .001 | 48, 8, 32 | 10.98 | < .001 |
| **aINS** | -28, 16, 8 | 12.41 | < .001 | 32, 20, 4 | 10.02 | < .001 |
| **IPS** | -40, -40, 44 | 10.52 | < .001 | 32, -54, 50 | 10.29 | < .001 |
| **Caudate** | -18, 2, 14 | 9.15 | < .001 | 18, 4, 16 | 8.98 | < .001 |
| **Putamen** | -22, -6, 6 | 11.87 | < .001 | 24, 0, 6 | 8.53 | < .001 |

Note: MNI coordinates and t-values refer to the local maxima within the clusters.

preSMA, pre-supplementary motor area; aINS, anterior insula; IPS, intraparietal sulcus.

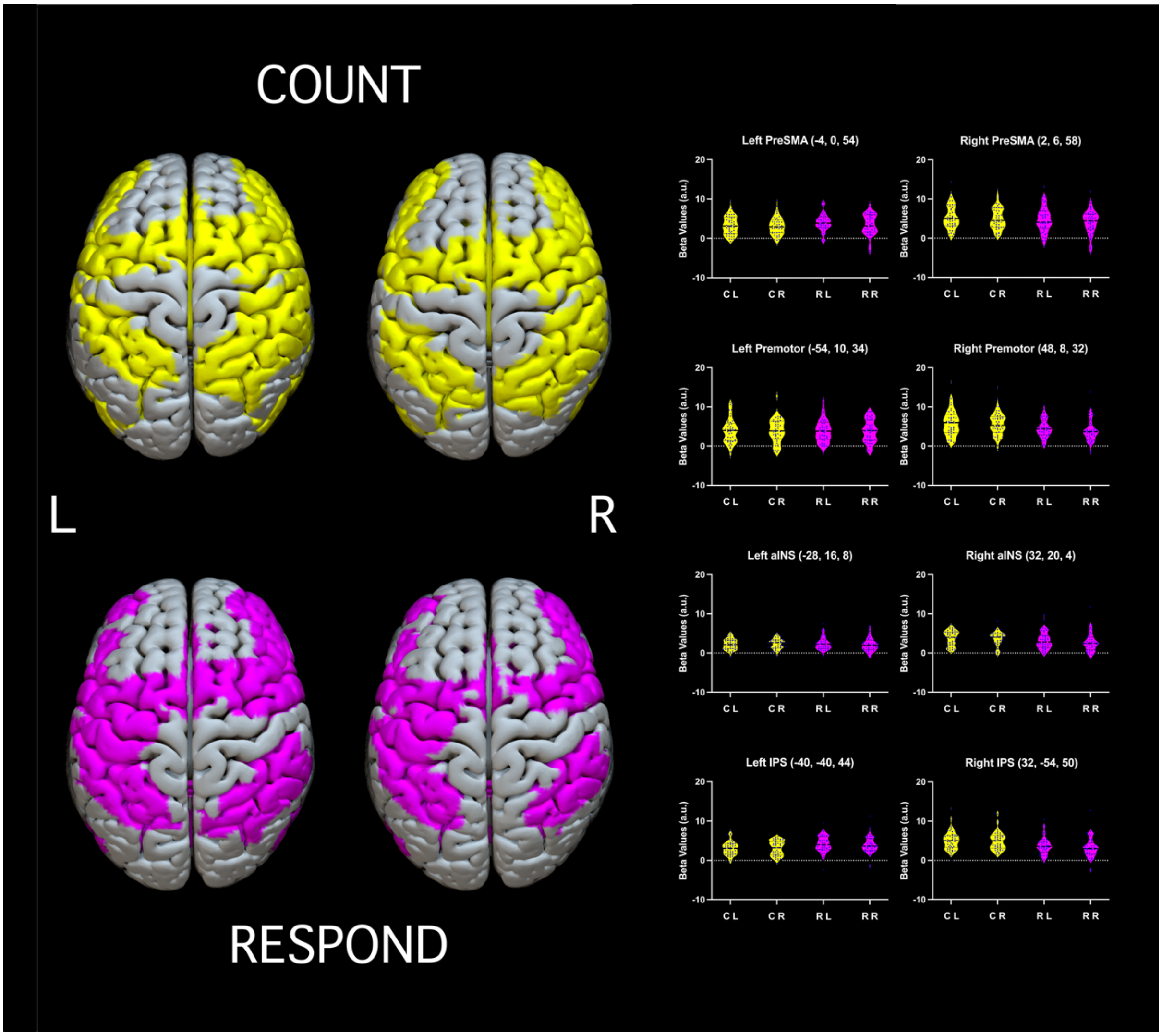


**Figure 1.** Thresholded images for Attend Motion Count Left and Attend Motion Count Right (yellow), and for Attend Motion Respond Left and Attend Motion Respond Right (magenta). Images shown are the voxelwise FWE-corrected ($p < .05$) contrast images used as input for the conjunction analysis.

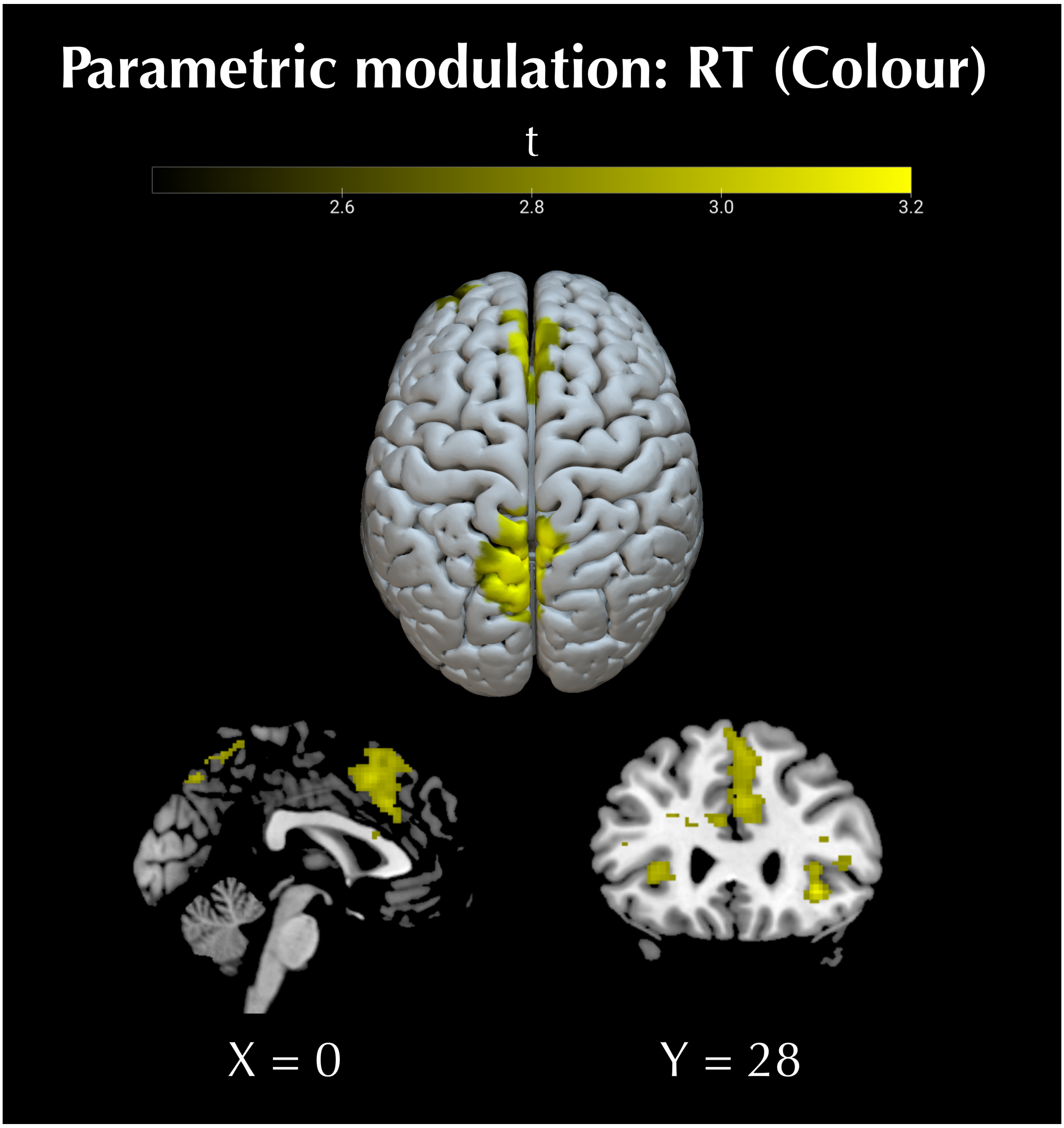


**Figure 2.** RT as a parametric modulator when attending to Colour. There were multiple significant clusters (*p* < .005, t > 2.71, FWE < .05 cluster-wise), including left superior parietal lobule (MNI: -6, -68, 52), bilateral anterior insula (MNI: 34, 28, -6), bilateral preSMA (MNI: -8, 8, 48), left ventromedial prefrontal cortex (MNI: -30, 46, 4), and left frontal operculum (MNI: -44, 18, 0).

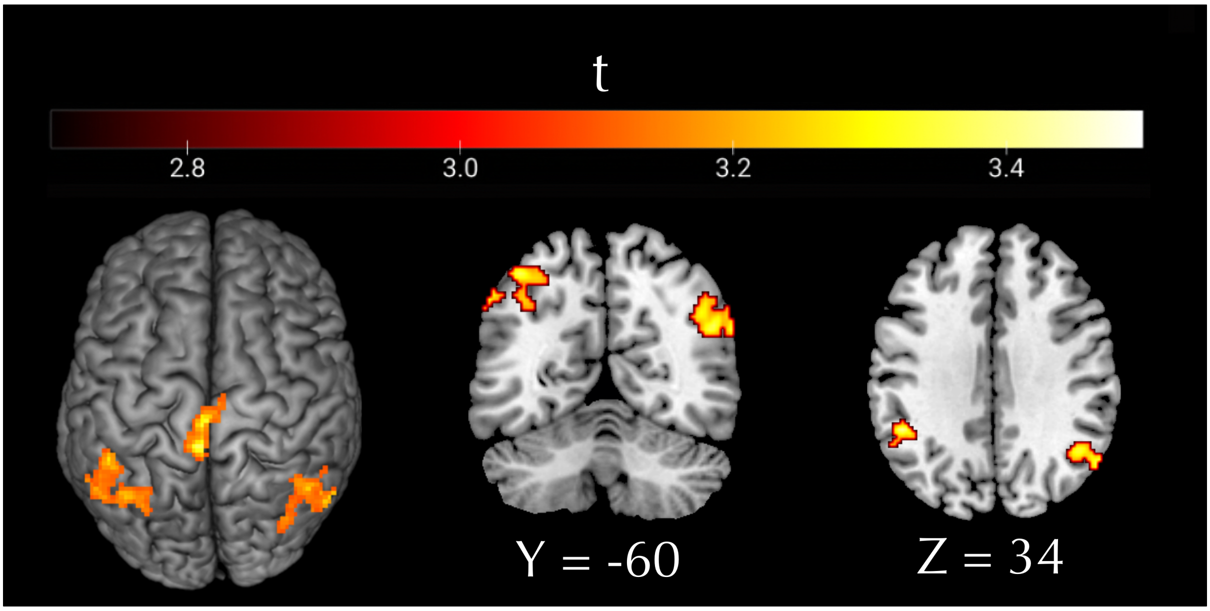


**Figure 3.** Whole-brain CPP slope regression. There were three significant clusters at the whole brain level ($p$ < .005, t > 2.71, FWE < .05 cluster-wise), the supplementary motor area (SMA) proper (MNI -4, -40, 70), bilateral inferior parietal lobule (MNI 50, -60, 30), and the left IPS (MNI -48, -46, 34). SMA activity has been linked to response preparation (Krainik et al., 2001; see Nachev et al., 2008 for review), and investigations of value-based decision-making have identified CPP activity within this region also (Pisauro et al., 2017).

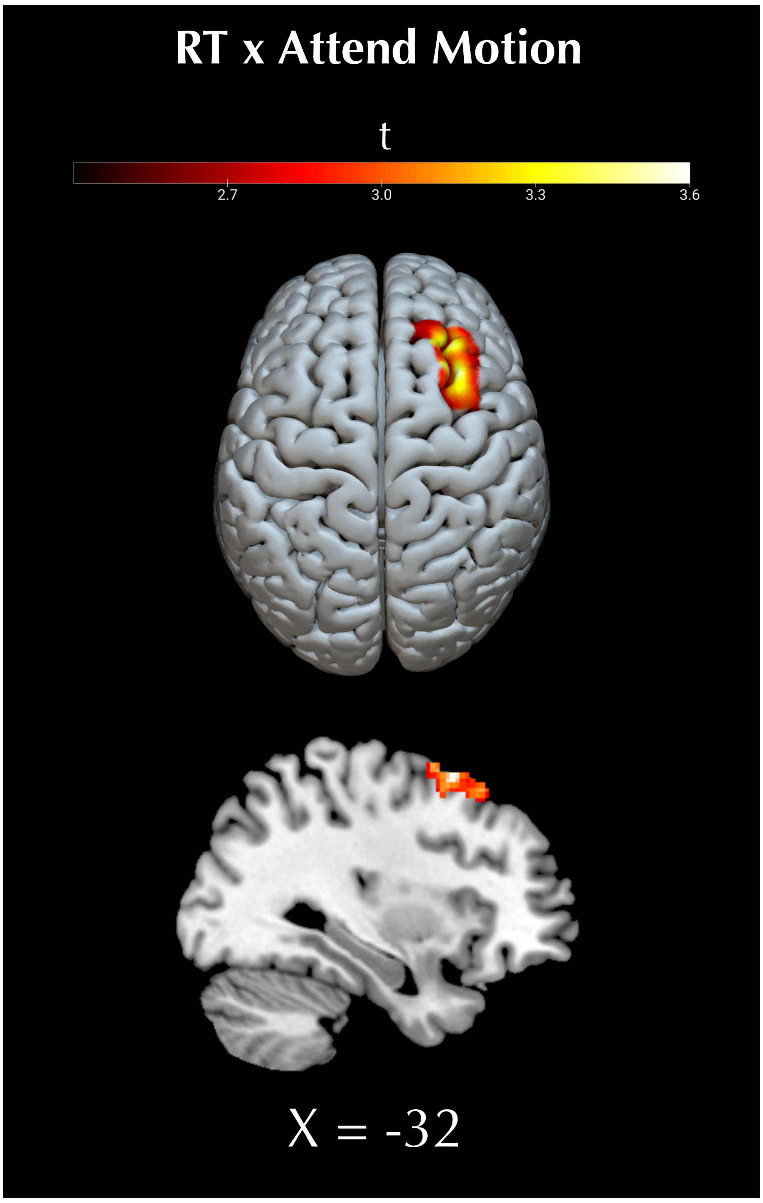


**Figure 4.** RT regression. There was a single significant cluster in the right middle frontal gyrus (MNI 32, 14, 60) ($p$ < .005, t > 2.71, FWE < .05 cluster-wise).

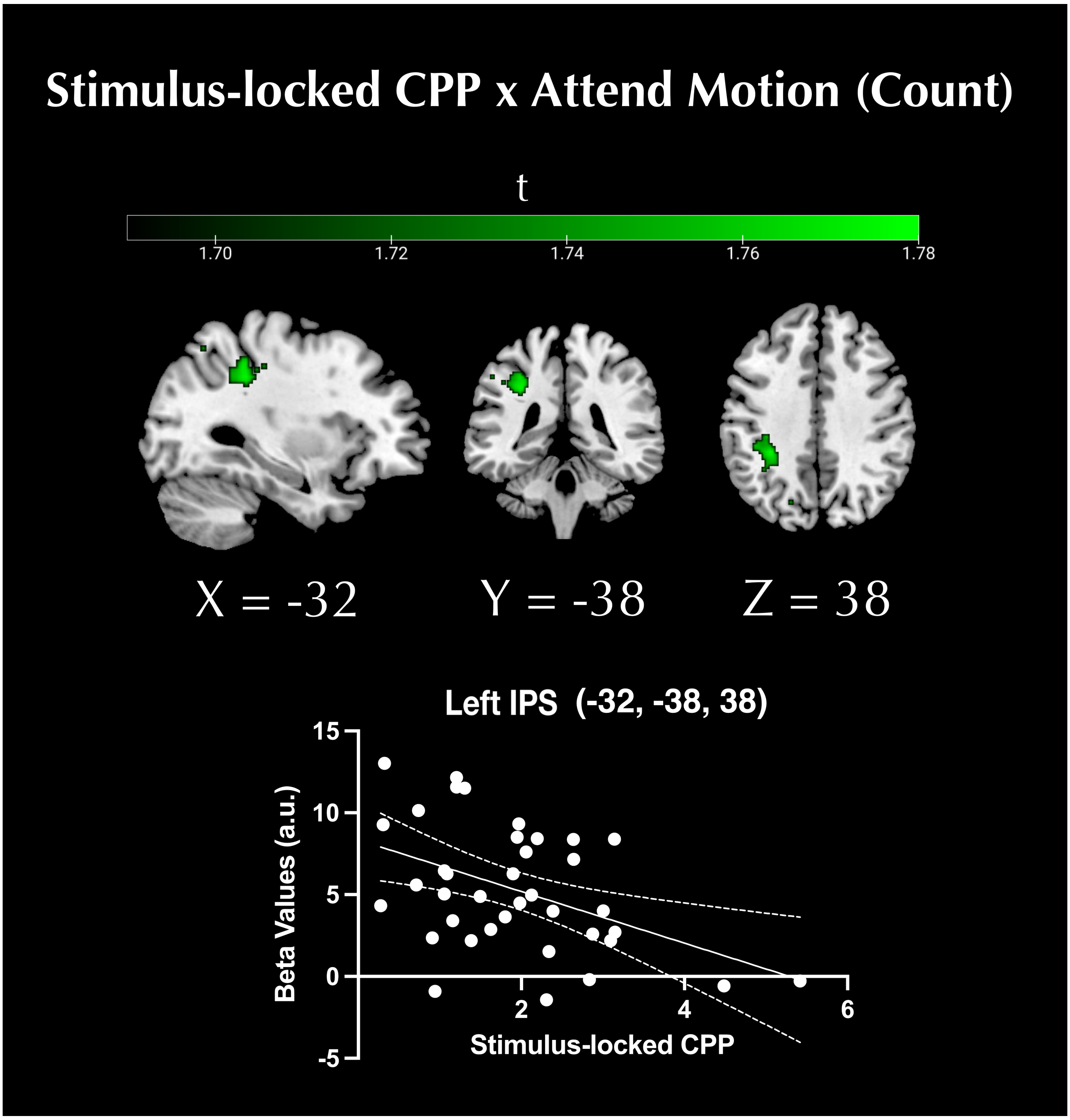


**Figure 5.** Stimulus-locked CPP slope as a regressor in a second level model using the Attend Motion Count > Rest contrast images for each participant. In the count condition there was no immediate response, and therefore it was not possible to derive a response-locked CPP. Instead, we could extract a stimulus-locked CPP, although the slope of the stimulus-locked CPP, like all stimulus-locked ERPs, contains substantial amounts of noise, including concurrent neural signals (McWeeny & Norton, 2020). Based on the accumulation-to-bound nature of the CPP (O'Connell et al., 2012) and established models of decision-making (Ratcliff & McKoon, 2008), the CPP (evidence accumulation) peaks at the time of response. Hence, although stimulus-locked CPP signals would still reflect evidence accumulation, the response-locked CPP is likely less noisy in comparison, rendering it more suitable for isolating the CPP's anatomical locus. Using the bilateral hIP3 ROI as an inclusive mask, and a liberal height threshold $p < .05$, $t > 1.69$, uncorrected, we see some evidence of a relationship between Attend to Motion (Count) BOLD activity and the stimulus-locked CPP slope in left anterior IPS.

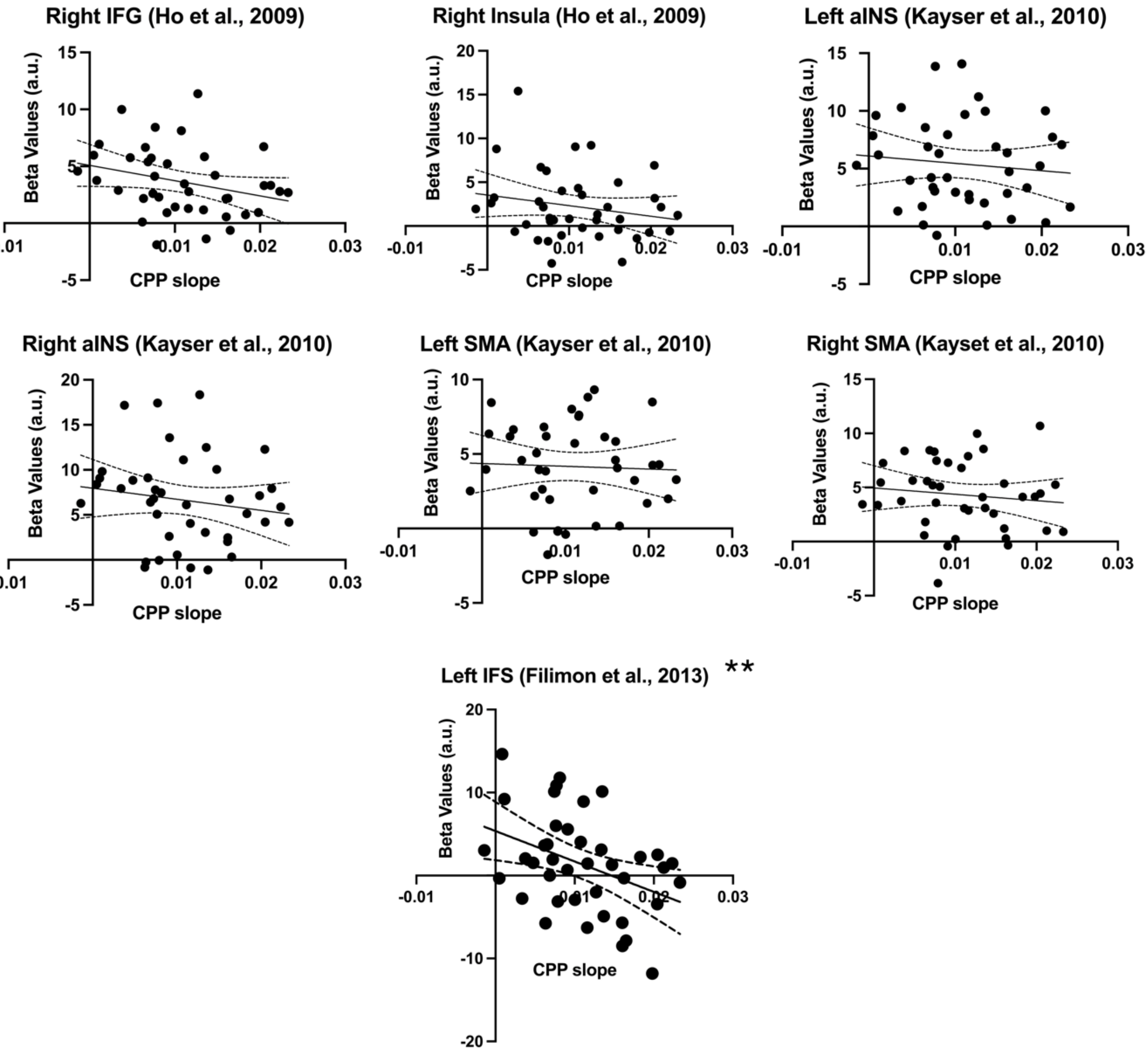


**Figure 6.** Examination of previously reported accumulator regions. Spherical ROIs (10mm radius) were defined around MNI coordinate peaks from previous studies and the extracted beta values were correlated with CPP slope.
Ho et al. (2009): Right IFG (MNI: 31, 17, 9), Right insula (MNI: 41, 7, 5);
Kayser et al. (2010): Left aINS (MNI: -32, 20, 9), Right aINS (MNI: 32, 23, 1), Left SMA (MNI: -10, 8, 47), Right SMA (MNI: 10, 14, 44);
Filimon et al., (2013): Left IFS (MNI: -48, 28, 22).
Note, IFG, inferior frontal gyrus; aINS, anterior insula; IFS, inferior frontal sulcus.
All correlations were non-significant ($p > .05$), with the exception of the left inferior frontal sulcus from Filimon et al. (2013) ($r = -.4$, $p = 0.01$).

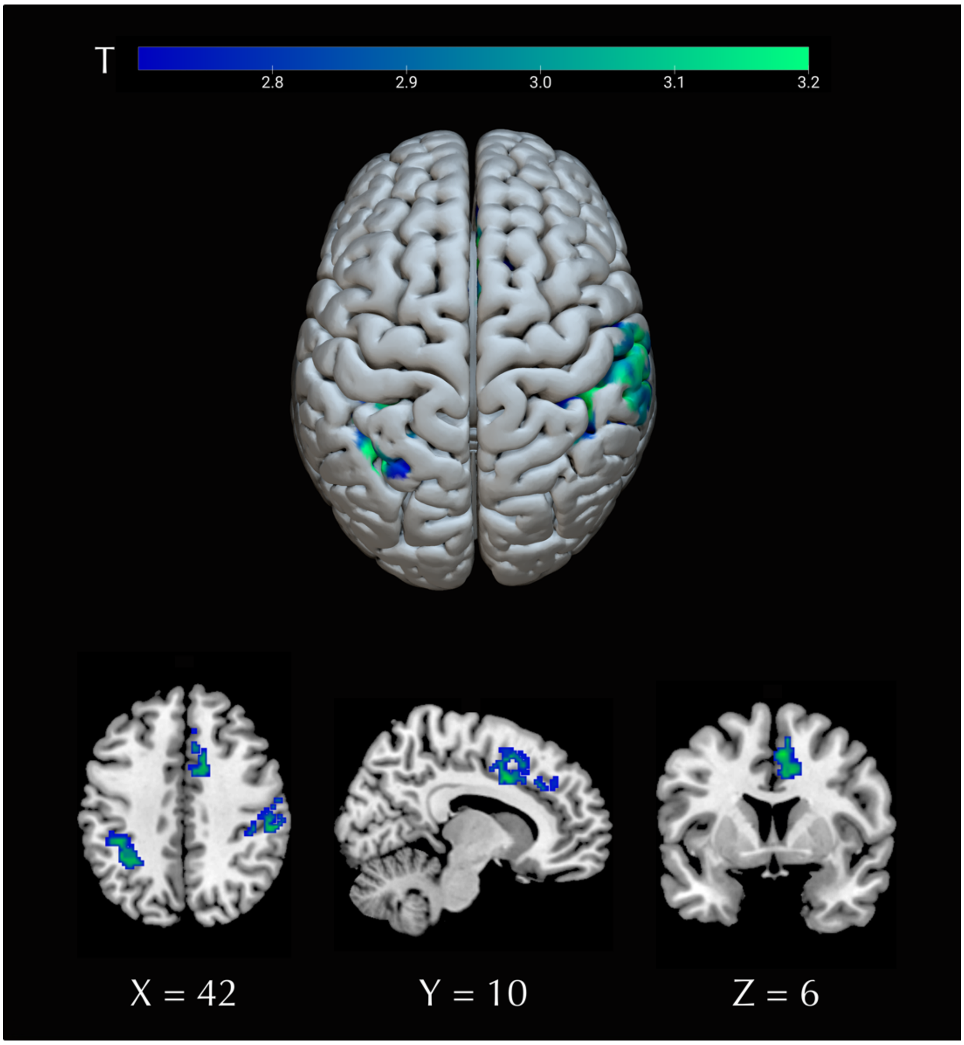


**Figure 7.** Whole brain analysis of drift rate as a parametric modulator. Images were set at a height threshold of $p < .005$, $t > 2.69$, FWE < .05 cluster-wise. Clusters were identified within left IPS (MNI -36, -50, 50), medial frontal cortex spanning the preSMA (MNI 6, 6, 48) and dorsal anterior cingulate cortex (MNI 12, 14, 38), and right inferior parietal lobule (56, -28, 42).

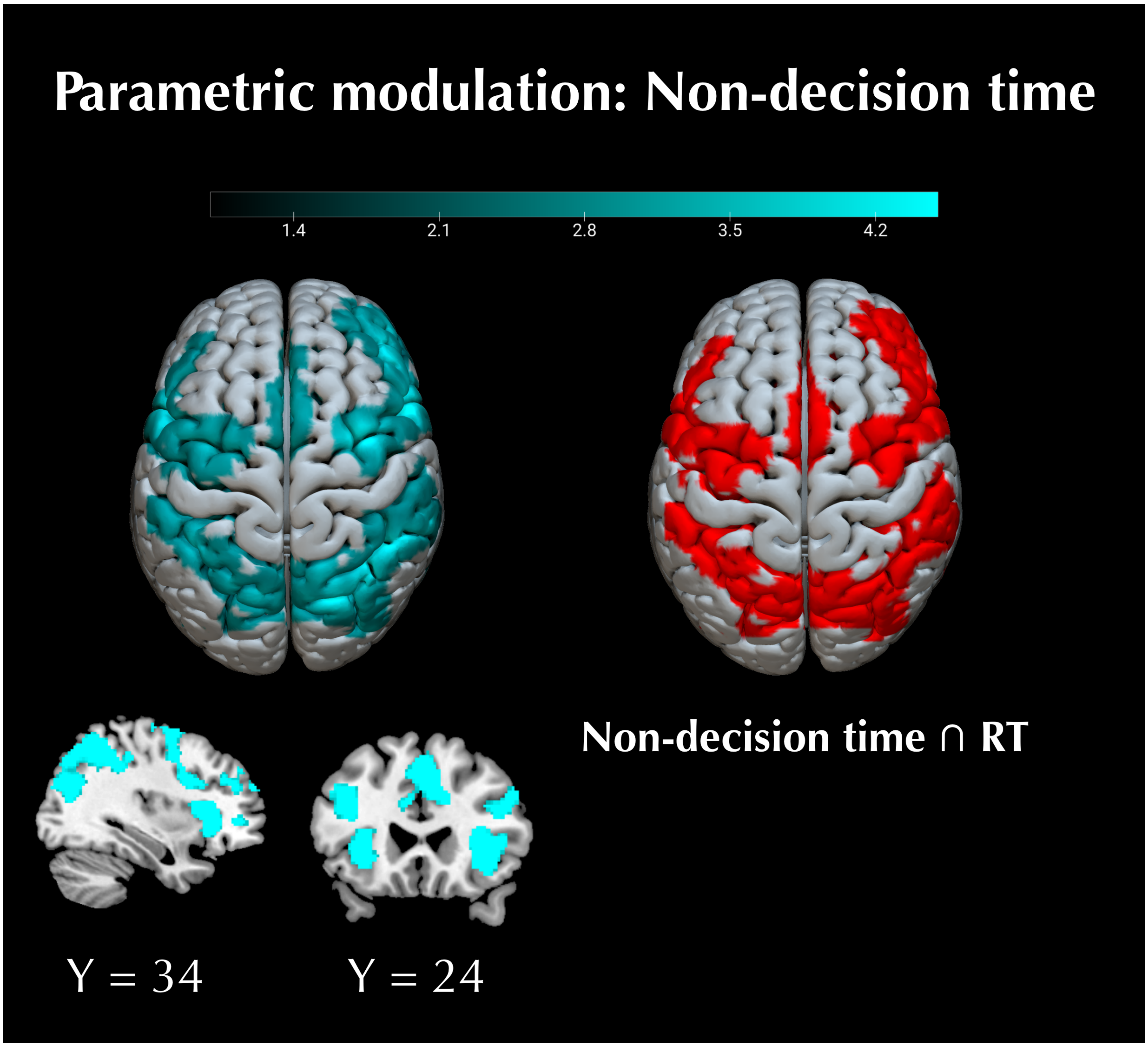


**Figure 8.** Whole brain analysis of non-decision time as a parametric modulator. Images were set at a height threshold of $p < .005$, $t > 2.69$, FWE < .05 cluster-wise. The conjunction of non-decision time and RT represents voxels that were common to both analyses.

**Supplementary Discussion**

***Colour onset***

Although we attempted to mitigate participants using the change in colour to prime participants' attention and thereby bias their perception, results revealed that Colour onset had an effect on RT when participants attended to Motion. Specifically, when Colour onset occurred after Motion onset, participants responded slower than when Colour occurred before, Mean RT difference = +29ms, SD = 4ms, $t(44) = 4.828$, $p < .001$; or coincident with Motion onset, Mean RT difference = +24ms, SD = 4ms, $t(44) = 3.727$, $p < .001$.

***Stimulus-locked CPP***

Notably, there was evidence of a relationship between CPP and left IPS when BOLD activity was regressed against stimulus-locked CPP slope values (note that a response-locked CPP cannot be derived in the count condition). However, the stimulus-locked CPP slope regression produced a more anterior IPS activation than the response-locked regression (Supplementary Data, Figure 5), which could suggest functional independence from the cluster identified using the response-locked CPP regression. It is possible that this is indicative of the fact that different parts of the IPS are responsible for evidence accumulation when motor responses are not required, which is consistent with some earlier research (Kayser et al., 2010; Filimon et al., 2013). Alternatively, this could reflect a limitation of using stimulus-locked CPP values given our low saliency stimuli. Smaller stimulus-locked CPP amplitudes have been associated with lower coherence stimuli (Kelly & O'Connell, 2015), which is consistent with the idea that using such stimuli may have resulted in weaker associations with BOLD activity.
